# Spatial summation of pain increases logarithmically

**DOI:** 10.1101/2020.06.30.179556

**Authors:** Wacław M. Adamczyk, Linn Manthey, Christin Domeier, Tibor M. Szikszay, Kerstin Luedtke

## Abstract

Pain intensity is difficult to predict. Mostly, because of modulatory processes underlying its formation. For example, when nociceptive stimulation occupies a larger body area, pain increases disproportionally. This modulation is called spatial summation of pain (SSp) and is responsible for coding pain intensity. To predict pain based on spatial variables, a profound understanding of the SSp effect is crucial. The aim of this study was i) to describe the SSp effect as a function of the size (or distance) of a stimulated area(s), ii) to investigate the effect of pain intensity on SSp and iii) to evaluate the influence of the SS type on the magnitude of SSp. Thirty-one healthy participants took part in a within-subject experiment. Participants were exposed to area- and distanced based SSp. In the former, electrocutaneous noxious stimuli were applied by up to 5 electrodes (5 areas) forming a line-like pattern at the ulnar side of the hand, while in the latter the same position and lengths of stimuli were used but only two electrodes were stimulated (5 separations). Each paradigm was repeated using pain of low, moderate and high intensity in a random and counterbalanced order. Each stimulus was assessed on a 0-100 scale. It was found that the pattern of increase in pain followed a logarithmic rather than a linear function. The dynamics of the pain increase were statistically different across pain intensities, with more summation occurring, if stimuli were calibrated to eliciting “high” pain. SSp was resistant to saturation in the area-based but not in the distance-based SSp, where 0.8cm separation between two electrodes produced a similar pain intensity as 1.6cm and 2.4cm. Results indicate that area-based SSp is more painful than distance-based SSp when low and moderate but not when high pain intensity is induced. Presented findings have important implications for all studies, in which the spatial dimension of pain is measured. When the area or separation between nociceptive stimulation increases, pain does not increase linearly. Furthermore, the pattern of the pain increase depends on i) intensity and ii) the number of sites of nociception. In conclusion, a logarithmic function should be considered when predicting the size of a nociceptive source. This pattern is indicative for inhibitory processes underlying SSp.

## INTRODUCTION

Pain is an adaptive response to noxious stimuli that protects humans from tissue damage. As in other sensory systems, nociceptive stimuli, which often -yet not always-lead to pain perception, are influenced by ascending (bottom-up) [56] and descending (top-down) [6,28,29] pain modulation. The same stimulus might be perceived as less or more painful depending on psychological, social and biological factors (see [1,14] for review). The latter category is particularly interesting as it reflects competitions between noxious stimuli targeting the neuroaxis with different temporal and spatial relations. For instance, research has shown that one stimulus is perceived as less painful if it is preceded by a heterotopic counter-conditioning stimulus [44,53], a phenomenon known as diffuse-noxious inhibitory-control (DNIC). However, if a pair of (or more) noxious stimuli occur concomitantly, they evoke pain facilitation, known as spatial summation of pain (SSp) [11–13,23,38,40,43,49,54].

Spatial summation of pain^1^ has been studied in animals [6,15,45] and humans [11–13,23,38,40,43,49,54] using two distinct paradigms: ‘distance-based’ and ‘area-based’ SSp [43]. In the former, an increase in pain is reported when the distance between stimuli increases, while in the latter, more pain is reported when noxious stimuli are applied to a larger area. Spatial summation of pain is essential for the detection of pain [2], coding of pain intensity [7] and identification of pain quality [9].

Transferring this knowledge to a patient population, the most obvious examples are patients with chronic widespread pain (CWP). Research in this population has shown that the intensity of pain perception can be predicted by the number of painful body parts [48,50], therefore, it is likely that the SSp mechanism is altered in this clinical population. Nevertheless, previous attempts failed to show differences in SSp response profiles between healthy controls and chronic pain patients [16,19,47,49], which raised the question about the mechanism of SSp. Interestingly, previous studies in which other paradigms, also testing pain modulation, such as temporal summation [34], offset analgesia [51] or DNIC, showed distinct differences between chronic patients compared to healthy controls.

The reason for this discrepancy is a relatively poor understanding of SSp in humans. Previous work has been mostly limited to psychophysical testing of SSp, with some physical or body-related manipulations made to study SSp mechanisms, e.g. skin types [17], modality [24,32], age [27], pain [49], stimulus intensity [42], body location [41], sequence of testing, [32] or the type of paradigm used [40] were considered. Notwithstanding to that, it is unclear where in the humans’ neuroaxis SSp occurs and how intensity shapes the magnitude of this effect. Behavioral studies are a first line of research that may approximate the underlying mechanisms of complex noxious processing.

In the past, SSp has been studied using rather ‘static’ paradigms. In that sense, area-based SSp was provoked by comparing responses to a small and a large probe. For distance-based SSp, various separations have been used to investigate this effect, often with small inter-spaced resolution, e.g. 4cm [40] 5cm [8,23,43] or 10cm [11] steps (separations). Whether summation also occurs using smaller scales (<1cm), has never been confirmed. Considering mentioned limitations, a mathematical modeling of the SSp response was not possible, due to the usually limited number of probes that were used.

The aim of this study was therefore to introduce a paradigm which investigates SSp as a function of a stimulated area to create adequate pain predictions. Secondly, the experiment aimed to investigate the effect of the experimentally induced pain intensity on the magnitude of SSp, described as a mathematical function. It was hypothesized that pain increases less dynamically as the pain intensity increases. Such a prediction is based on the evidence showing that more pronounced pain inhibition occurs, when more severe [46,55] and larger [24,32] stimuli are used. The associated higher activation of pain-inhibitory brain regions is linked to stimulus aversiveness. Lastly, we developed a study design to investigate SSp at a micro-scale level, with stimuli forming a continuum of < less than 4cm in length. Such a design allows to explore the relationship between the type of SSp (area-based, distance-based) and the effect of the stimulus intensity, as well as their interactions.

## MATERIALS AND METHODS

### Study overview and structure

This study was based on a within-subject design to test the effect of SSp type and stimulus intensity on the pattern of pain summation. The structure of single session was similar to those described previously [33,39] and was subdivided into familiarization, calibration and the main data collection phase (SSp assessment). During the last phase, participants were assessed using two paradigms (area- and distance-based SSp) with three different pain intensities (low, moderate and high pain). Three sessions were performed in total.

The Ethics Committee of the University of Lübeck approved the protocol of this study (decision no. 19-303), which was preregistered in December 2019 at the osf.io platform (https://osf.io/qry9d), using the AsPredicted.org template. The study follows the principles of the Declaration of Helsinki, developed by the World Medical Association. Each participant was adequately informed about the objectives, methods, the anticipated benefits and potential risks and the discomfort, as well as any other relevant aspects of the study. A written informed consent was obtained from each participant before participation in the study.

### Study sample

Fifty-three volunteers signed up to participate in the study. Each volunteer was screened for eligibility. To be included in the study, participants had to be healthy (self-report), right handed, over the age of 18 years and have sufficient German or English language skills. Exclusion criteria were any acute or chronic pain, skin pathologies or tattoos in the area of the left hand, diagnosed neurological, cardiovascular or psychiatric diseases or any other disease requiring regular medication intake. Other exclusion criteria were pregnancy, metal implants or electronic devices in or on the body. The exclusion criteria were established to avoid any changes in pain perception and possible health-related risks during the application of electronically induced pain. Twenty-two volunteers were not considered eligible and 31 (15 females, 48.39%) participated and completed the experiment (mean age 26.2 ± 6.8years, height 174.0 ± 9.9cm, weight 68.9 ± 13.5kg).

As it has been shown that both, caffeine and alcohol consumption, affect the sensation of pain [25,52], participants were also asked to refrain from consumption on the day of participation. They were further asked to not take any drugs 24 hours before participating in the study. In order to avoid a DNIC effect, participants were asked to not engage in excessive sports that could cause muscle ache or other pain 48 hours before participation in the study.

### Sample size

Previous studies investigated SSp in healthy subjects using sample sizes of 25 [43] or 20 [23] participants, however the effect of intensity has not been tested in those studies. To avoid underestimation of the effects of the stimulation area and the intensity used, it was decided to assess at least 30 participants. Such a number of participants should suffice to test for a moderate effect size with 80% power and *α* = 0.05, as reported previously [23,43].

### Experimental setting and materials

The trial took place in a quiet and temperature-controlled (20.5 ± 0.5 °C) laboratory. The participants were set on a chair in front of a monitor with a distance of 50cm. The 23.8-inch monitor was set up 1m above the floor and the chair was 50cm high. The participant was separated from the examiner by a moveable curtain. Pain was assessed using a Numerical Rating Scale (NRS) after each stimulus application. The scale was anchored from 0 (“no pain”) to 100 (“most imaginable pain”) [43].

The electrocutaneous stimuli were produced by a Constant Current Stimulator (Digitimer, model DS7A, United Kingdom, Welwyn) and a remote electrode selector activated a given set of electrode(s) (Digitimer, model D188, United Kingdom, Welwyn) with 200µs duration of each electrocutaneous pulse and a capacity of 0 to 100 mA with a maximum voltage of 400V, which is based on previous SSp studies [23,43]. One stimulus consisted of a train of 20 (square) pulses (inter-pulse interval 10ms). External control of DS7A and D188 was ensured via the Labjack U3-LV control device (LabJack Corporation, Lakewood, CO, USA). The procedure was fully automatic and operated by the PsychoPy 3.0, open-source software [36]. Five 8-mm diameter, planar concentric, electrodes (WASP electrodes, Brainbox Ltd., Cardiff, UK) were used to stimulate nociceptive fibers. Electrodes consisted of two gold plated solder pads, with a platinum cathode in the centre and a concentric anode [37]. They were attached first to the armrest of the armchair, where the participants were seated and later to their hand. Before attaching the hand electrodes, the ulnar edge of the hand was cleaned with an abrasive gel and an alcohol wipe using a cotton pad to remove all particles, which could have a negative impact on the skin impedance.

Electrodes were placed just below the skin fold under the metacarpophalangeal joint of the fifth phalanx (ulnar side of hand). The hand was placed in a neutral position between pronation and supination and the elbow was rested at 90° of flexion. Electrodes were placed next to each other (with no space between adjacent electrodes). Such an orientation allowed to study area-based SSp with up to 5 electrodes activated simultaneously. For distance-based SSp, the same configuration was used. However, distanced-based SSp was assessed when two electrodes (with randomly chosen distance between them), were activated. In principle, area-based SSp is established when one manipulates the size of the stimulated area and distance-based SSp, when different separations are used. Having this conceptual difference in mind, the following definitions were used for clarity: Different areas were determined by the number of electrodes activated (up to 5) and different separations were determined by the number of electrodes forming the gap between two activated electrodes. As the electrodes were placed in a line, activation of 5 electrodes induced a stimulus of 4cm length (5 *×* 0.8cm) that served as the maximal area in area-based paradigm. The same distance (4cm), formed by two outermost electrodes, was used as the maximal separation for the distance-based paradigm (Fig. 2). Thus, technically, the stimuli had the same length, regardless of the SSp type being assessed.

**Figure 1.**
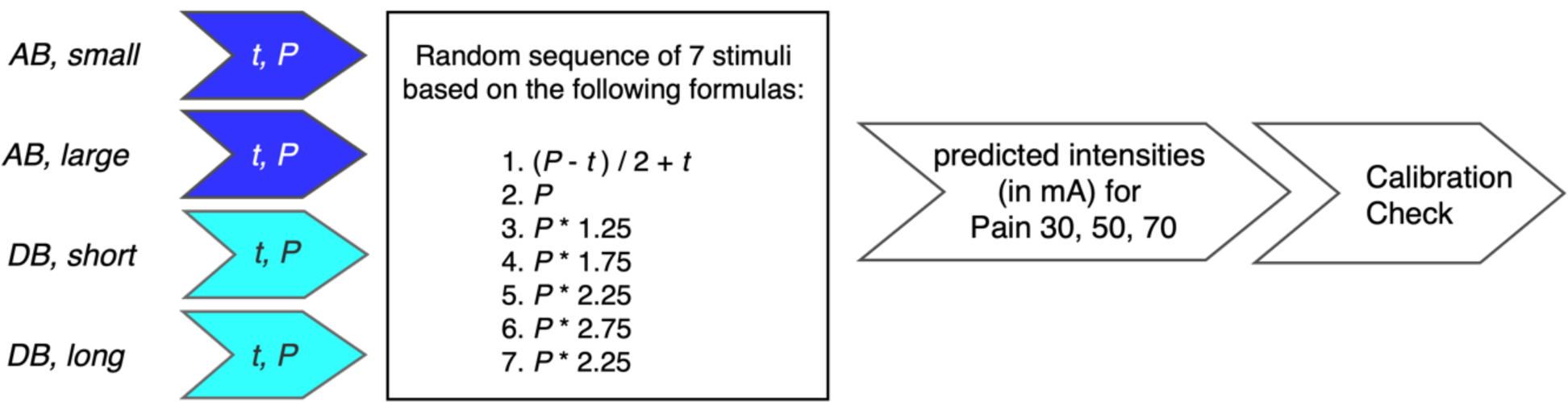
Calibration phase. Calibration started from tactile (t) and pain threshold (P) determination using different electrode orientations. Based on these results, 7 stimuli of different intensities were applied in a random order. Then different pain intensities corresponding to low (30 on Numeric Rating Scale, NRS), moderate (50) and high (70) pain were readout from regression lines plotted as a function of pain and intensity (in mA). In the end, the given intensity was checked for accuracy in the pain induction of assumed intensity (calibration check).

**Figure 2.**
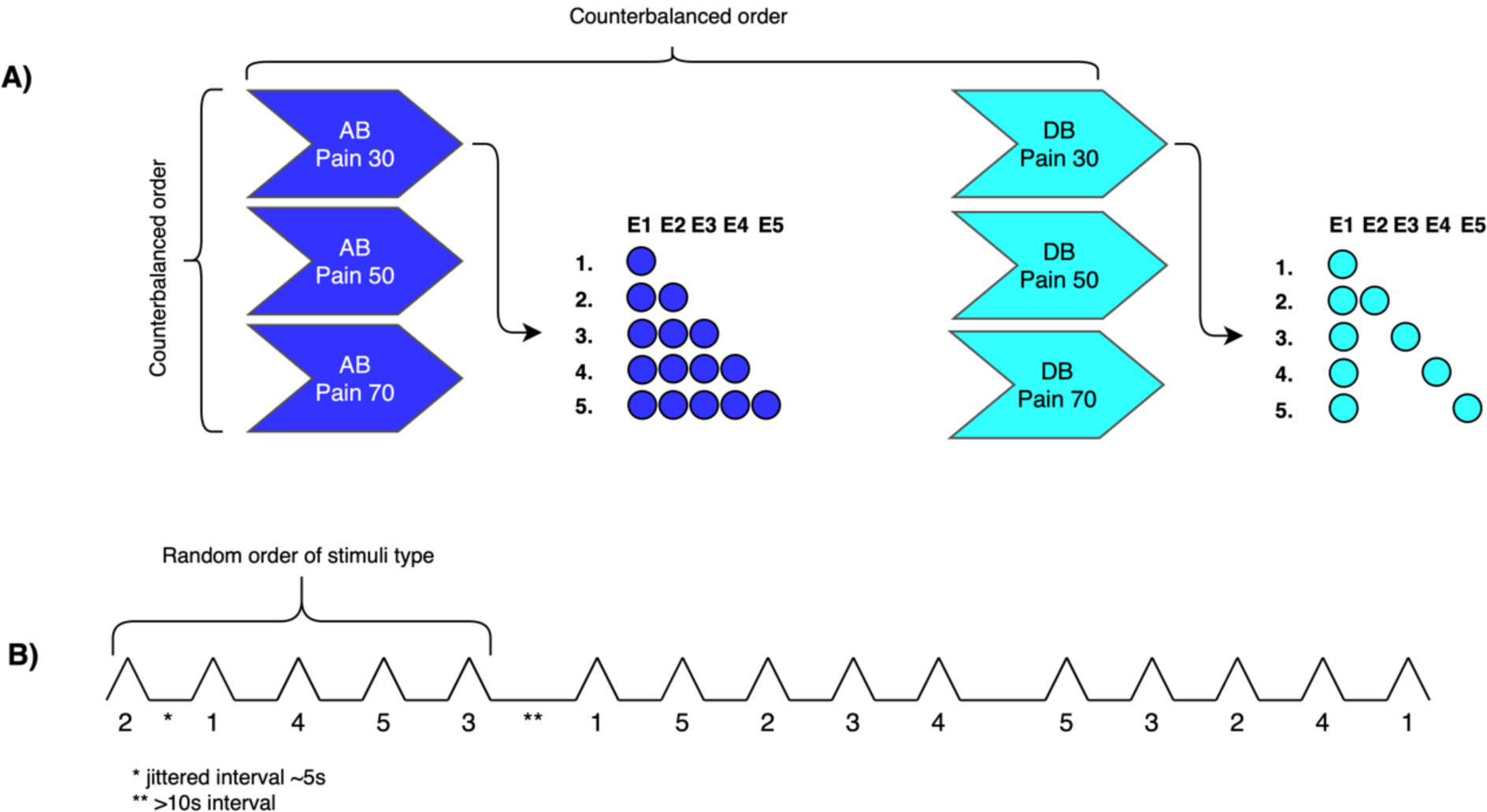
Structure of the spatial summation assessment (SSp). **A)** For every pain condition (pain at the level of 30, 50, or 70 on 0-100 NRS scale), 5 different stimuli configurations were used. In the area-based (dark blue) SSp either one electrode or up to 5 electrodes were activated simultaneously. In the distance-based paradigm (cyan) either one electrode or two electrodes with a varied separation were activated simultaneously. In principle, for both SSp types, stimuli formed line-like pattern, with filled or unfilled areas (with different separations). The order of the pain conditions and electrode configurations were randomized and counterbalanced across participants. **B)** For every pain condition there were three series consisting of 5 stimuli each. In total, 15 stimuli were applied for every pain intensity, thus 45 stimuli were used per SSp type (90 stimuli in total). The order of the stimuli was randomized and counterbalanced across the assessment. AB, area-based SSp, DB, distance-based SSp, E, electrode. Note, that the same intensities determined individually for the given condition (low, moderate and high pain), were used for both SSp types.

### Familiarization and calibration

To check if the conduction under all 5 electrodes is accurate, an initial testing procedure was performed. The electrodes were activated sequentially with a fixed intensity of 6.0mA and inter stimulus intervals (ISI) of 5s. Detection of the stimuli were verbally reported by the participants after each trial. This procedure was also performed to make participants familiar with electrocutaneous stimulation.

### Calibration

Participants underwent 4 series of calibration trials to consider the individual differences in pain perception and to determine the individual intensities for the subsequent assessment of SSp. The objective of the calibration was to obtain pain corresponding to an NRS of 30 (low), 50 (moderate) and 70 (high intensity). Thus, the influence of different pain conditions on the magnitude of SSp could be evaluated.

To calibrate the individual intensity levels, first the tactile (*t*) and pain (*P*) thresholds were determined [4,5]. This was done with electrode 1 and 2 activated with a separation of 0cm (shortest possible distance) for distance-based SSp and also with only one electrode activated (smallest possible area) for the area-based SSp. Furthermore, *t* and *P* were determined for the largest distance possible, which is equivalent to an activation of electrode 1 and 5 (distance-based SSp) and also with the largest area, which is equivalent to an activation of all 5 electrodes (area-based SSp). For each calibration part, the electrical current started from 0.0mA and was increased by 0.5mA, until the participant verbally reported a very first sensation (*t*). The intensity was further increased by 0.5mA until the first pain perception was reported (*P*).

Based on these determined thresholds, participants received a random sequence of painful and non-painful electrocutaneous stimuli of different intensities. The intensities were calculated using the formulas depicted in Fig. 1. The participant was instructed to rate every stimulation on the NRS by selecting a level that best corresponded to the perceived intensity. Based on these pain ratings, a regression line for the stimulus-response function was individually plotted. The x-axis represented the electrical current in mA, the y-axis the individual pain ratings.

Through the formula obtained from the given regression line, a predicted intensity (in mA) for all pain conditions (NRS 30, 50 and 70) was determined, using small (short) and large (long) configurations (Fig 1). An average of predicted electrical currents was then tested in the calibration check [31]. The calculated stimulus intensities for the different pain levels were applied during the check, if predicted values from the regression models induced pain at the targeted level (pain of 30, 50 and 70, respectively). The same final intensities were used for both SSp types, to compare pain summation between paradigms.

### Spatial summation assessment

To assess SSp, different stimuli configurations were used, as described earlier. The line-like pattern of 5 electrodes, allowed to apply 5 different stimulus intensities within each SSp type (see Fig. 2). For the area-based paradigm, the control (one) electrode was activated separately or simultaneously with a sum of up to 5 electrodes. Therefore, stimuli forming a filled line ranging from 0.80cm in length up to 4cm were applied. A series of 5 stimuli with jittered 5-7s ISI was applied in a fully random sequence and repeated three times (>10s break between the series) for each pain condition, i.e. low, moderate and high pain.

For the distance-based paradigm the same structure was maintained. Here the stimuli had the same line-like pattern (from 0.80 to 4cm), however, the control electrode was activated separately or simultaneously with a second electrode 0.0, 0.8, 1.6, or 2.4cm apart (Fig. 2). The order of SSp type, and the order of pain conditions (low, moderate, high) was randomly assigned, yet counterbalanced across subjects. For each paradigm (area- and distance-based SSp), there were 45 stimuli induced, representing 5 configurations *×* 3 series *×* 3 intensities, so that each participant received 90 stimuli in total during the assessment phase. As there were 3 sessions employed, each participant received 270 stimuli.

### Data processing and statistical analysis

Raw pain ratings were first extracted from the files generated by the PsychoPy software using MATLAB R2017b (MathWorks Inc, Natick, Massachusetts). Descriptive statistics of all collected data were presented as means and SD or their non-parametric equivalents. The analysis was based on a complete dataset from 31 participants.

Spatial summation of pain was analysed in according to the following protocol: To reduce data for the main analysis, a General Linear Model (GLM) was applied to test for significant difference in pain perception between measurement sessions. Next, the main SSp analyses were performed on the pooled dataset from three sessions. Namely, a 2 *×* 3 *×* 5 GLM analysis was performed with within-subject factors ‘paradigm’ (area-based, distance-based SSp), ‘intensity’ (low, moderate, high) and ‘stimulus configuration’ (line of 0.8, 1.6, 2.4, 3.2, 4cm length). In case of significant main or interaction effects, post-hoc Bonferroni corrected Tukey tests were applied to investigate the meaning of the effect.

To express SSp in one unified coefficient, linear (*y* = *b*\**x* + *a*) and logarithmic (*y* = *b*\*log(*x*) + *a*) curves were fitted to individual subjects for area-based and distance-based SSp in respect to stimulus intensities. Thus, six linear and logarithmic curves were fitted to the data per subject using the Curve Fitting Toolbox in MATLAB R2017b (MathWorks Inc, Natick, Massachusetts). Goodness of fit was assessed visually and by comparing R^2^ statistics generated by two different fittings using paired t-tests and by one-sample t-tests against perfect fit of 1.0 *R*^*2*^ value. The magnitude of SSp was expressed as the slope (*β*-coefficient) of the curve where -in case of logarithmic distribution of the data-higher values mean more pronounced SSp as the summation increases more dynamically together with the linear increase in the stimulated area (or separation). To further demonstrate the SSp effect across different intensities, a GLM model was used with *β*-coefficients as dependent variables and ‘paradigm’ (area-based and distance-based SSp) and intensity (low, moderate, high) as two within-subject factors. Post-hoc Tukey tests were applied - if necessary - with Bonferroni correction to control for family-wise error.

Statistical analyses were conducted using the STATISTICA data analysis software, version 13 (StatSoft Inc., Tulsa, OK, USA). The level of significance was set at *p* < 0.05. When *p*-values did not exceed *α* levels after correction, it was marked as not significant (ns).

## RESULTS

Thirty-one healthy participants took part and finished this experiment. Participants’ characteristics are presented in Table 1. General Linear Model on ‘pain ratings’ as dependent variable and ‘session’ and ‘stimuli’ as within-subject factors revealed no significant effect of ‘session’, neither for area-based (*F*_(2,60)_ = 0.61, *p* = 0.54,*η*_*p*_^2^ = 0.02) nor for distance-based SSp (*F*_(2,60)_ = 1.16, *p* = 0.32, *η*_*p*_^2^ = 0.04). Thus, datasets were combined across sessions and subsequent analysis was performed on pooled dataset.

**Table 1.** Characteristics of study participants (n = 31)

| Variable | Mean | SD |
| --- | --- | --- |
| Age in years | 26.19 | 6.83 |
| Palmar-wrist crease in cm | 16.5 | 1.4 |
| Hypothenar length in cm | 6.58 | 0.93 |
| Weight in kg | 68.87 | 13.46 |
| Height in cm | 173.97 | 9.88 |
| Fear of pain | 3.23 | 1.82 |
| PVAQ | 39.39 | 9.14 |
| PCS | 9.58 | 8.16 |

SD, Standard Deviation, PVAQ, Pain Vigilance and Awareness Questionnaire, PCS, Pain Catastrophising Scale.

The main GLM analysis performed on pain ratings showed a significant main effect of ‘SSp’ (*F*_(1,30)_ = 21.47, *p* < 0.001,*η*_*p*_^2^ = 0.42), ‘intensity’ (*F*_(2,60)_ = 245.47, *p* < 0.001, *η*^2^*p* = 0.89) and ‘stimuli’ (*F*_(4,120)_ =122.50, *p* < 0.001, *η*_*p*_ = 0.80). These three main effects indicated that i) in general, area-based SSp was more painful compared to distance-based SSp (Fig. 3 and 4), ii) calibration was successful as participants discriminated pain levels significantly (*p* < 0.001), and iii) SSp was statistically significant as -in general-pain increased with the area or distance between two stimuli (*p* < 0.001, Fig. 3). Furthermore, the GLM showed significant ‘SSp’ *×* ‘intensity’ (*F*_(2,60)_ = 3.26, *p* < 0.05, *η*^2^_*p*_ = 0.10), ‘intensity’ *×* ‘stimuli’ (*F*_(8,240)_ = 9.19, *p* < 0.001, *η*^2^_*p*_ = 0.23) interactions as well as an interaction between the factors ‘SSp’ and ‘stimuli’ (*F*_(4,120)_ = 76.82, *p* < 0.001, *η*^2^_*p*_ = 0.72).

**Figure 3.**
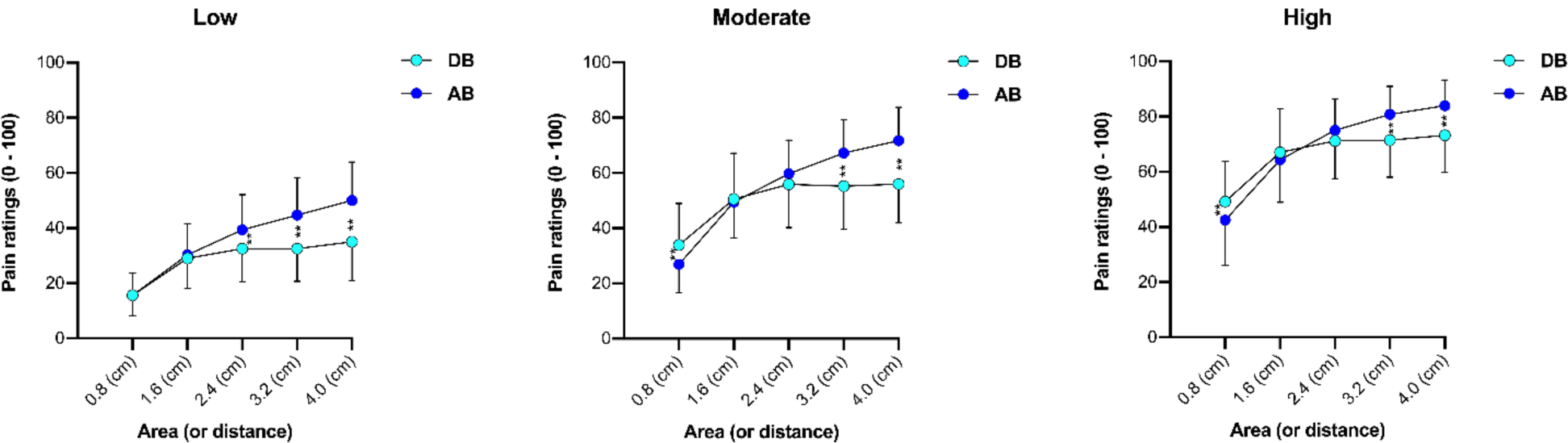
Spatial summation of pain (SSp). The pain intensity increases with increasing the stimulated area (AB) or distance (DB) between two stimuli in all pain conditions. Behaviour of area-based (AB) and distance-based (DB) SSp under the pain of low (left), moderate (middle) or high intensity (right). Note that in general, area-based SSp was more painful than distance-based but not when the highest intensity was used. Each figure presents mean values (dots) with standard errors of the mean (SE). ** *p* < 0.01.

**Figure 4.**
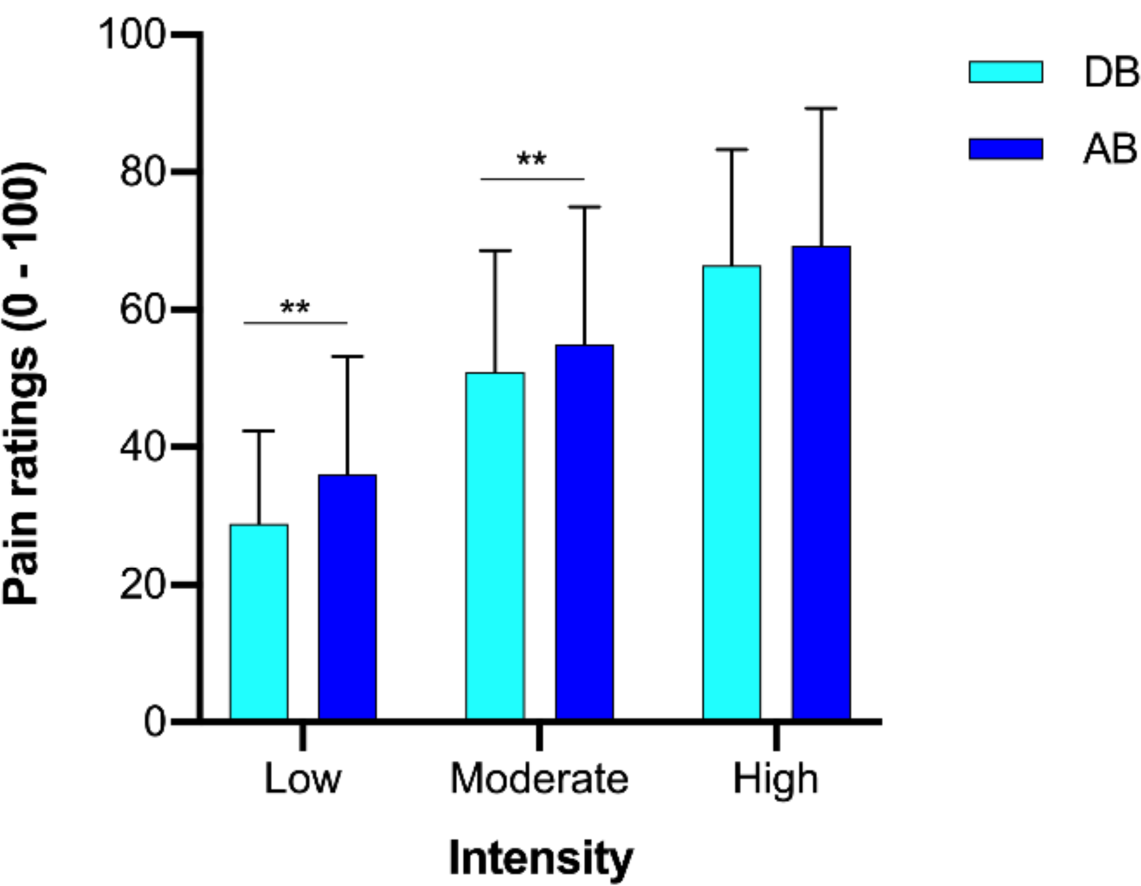
Pain intensity across spatial summation paradigms. Comparing the types of SSp, the area-based (AB) paradigm was perceived as more painful compared to the distance-based (DB) paradigm except for the highest intensity: here relatively the same level of pain was induced even though 2.5 times less area was stimulated in distance-based paradigm. ** *p* < 0.01.

Results of post-hoc tests following these interaction effects revealed that i) area-based was more painful than distance-based SSp but only when low (*p* < 0.001) or moderate (*p* < 0.001) but not when high intensity (*p* = 0.16) was applied (Fig. 4), ii) further increases in area or distance did not lead to a systematic increase in pain perception; regardless of the SSp type, the SSp effect was saturated when comparing the two outermost stimuli, i.e. 3.2cm vs. 4cm length (Fig. 3). The stimulation of the largest area consisting of 5 electrodes produced similar pain level compared to the slightly smaller area formed by 4 electrodes when low (*p* = 0.06), moderate (*p* = 0.10) or high (*p* = 0.18) intensity condition was considered, iii) the behaviour of SSp is different across SSp types: Area-based SSp was not saturated over the course of the linearly increased size of the stimulated area, as the pain level still increased significantly when the stimulus was enlarged (all *p* values < 0.01, except for 0.8 vs. 1.6cm stimuli - n.s. after correction). However, in the distance-based paradigm, pain was saturated and did not further increased when enlarging the distance, 0.8cm separation produced similar pain as 1.6cm (Fig. 3, *p* = 0.99) and 2.4cm (*p* = 0.77).

Lastly, GLM analysis showed a significant three-way ‘SSp’ *×* ‘intensity’ *×* ‘stimuli’ interaction (*F*_(8,240)_ = 3.06, *p* < 0.01,*η*^2^_*b*_ = 0.09), indicating that the pattern of pain increase was different across the SSp types in respect to the intensity. Similar pain levels across SSp types were observed when the stimulus of 1.6cm (or 0cm separation between 2 stimuli) was applied with low (*p* = 0.99) moderate (*p* = 1.00) and high (*p* = 0.43) intensity. Interestingly, starting from the third level of stimuli (2.4cm stimuli length), area-based was more painful than distance-based SSp (Fig. 3), however, when a single stimulus was contrasted across SSp paradigms, area-based SSp was paradoxically less painful for moderate (*p* < 0.05, Fig. 3) and high intensity (*p* < 0.05, Fig. 3). This, however, might reflect procedural aspects of the experimental paradigm, such as a higher pain intensity contrast between different stimuli in the area-based paradigm.

**Table 2.** Descriptive statistics for pain ratings in two SSp types

| Intensity | Area (separation) | Area-based |  | Distance-based |  |
| --- | --- | --- | --- | --- | --- |
|  |  | Mean | SD | Mean | SD |
| Low pain<br>(30/100) | 0.8 cm | 15.64 | 8.31 | 15.64 | 7.57 |
|  | 1.6 cm | 30.29 | 11.54 | 28.87 | 11.54 |
|  | 2.4 cm | 39.37 | 12.96 | 32.56 | 12.19 |
|  | 3.2 cm | 44.70 | 13.73 | 32.57 | 12.08 |
|  | 4.0 cm | 50.03 | 14.19 | 34.76 | 14.12 |
| Moderate pain<br>(50/100) | 0.8 cm | 26.84 | 10.37 | 33.84 | 15.34 |
|  | 1.6 cm | 49.48 | 13.23 | 50.52 | 16.89 |
|  | 2.4 cm | 59.67 | 12.44 | 55.86 | 15.92 |
|  | 3.2 cm | 67.20 | 12.17 | 55.20 | 15.90 |
|  | 4.0 cm | 71.70 | 12.30 | 55.95 | 14.27 |
| High pain<br>(70/100) | 0.8 cm | 42.48 | 16.74 | 49.24 | 14.88 |
|  | 1.6 cm | 64.35 | 15.63 | 67.08 | 16.05 |
|  | 2.4 cm | 75.02 | 11.50 | 71.22 | 14.07 |
|  | 3.2 cm | 80.76 | 10.40 | 71.48 | 13.63 |
|  | 4.0 cm | 83.94 | 9.53 | 73.18 | 13.67 |

SD, Standard Deviation.

Studying pain behaviour across the full range of areas and distances allowed for visual assessment of the stimulus-response functions (Fig. 3), which suggested that the pattern of pain increase followed logarithmic function. When comparing *R*^2^ coefficients for logarithmic and linear models by using one-sampled (*R*^2^ against referenced value of 1.0) and two sampled paired *t* -tests (*R*^2^ across models), it was found that i) lower *t* statistics were observed for the logarithmic compared to the linear fitting (Table 3) and ii) that logarithmic curves showed significantly higher *R*^*2*^ statistics compared to linear fittings (Table 4).

**Table 3.** Curve fitting: one-sample t-tests results (mean *R*^2^ value against value of 1)

| SSp | Intensity | N | Logarithmic |  |  |  | Linear |  |  |  |
| --- | --- | --- | --- | --- | --- | --- | --- | --- | --- | --- |
| | | | $R^2$ | SD | $t$ | $p$ | $R^2$ | SD | $t$ | $p$ |
| AB | Low | 31 | 0.86 | 0.11 | -7.40 | < 0.001 | 0.81 | 0.11 | -9.31 | < 0.001 |
|  | Moderate | 31 | 0.86 | 0.09 | -8.64 | < 0.001 | 0.80 | 0.12 | -9.74 | < 0.001 |
|  | High | 31 | 0.88 | 0.07 | -9.52 | < 0.001 | 0.79 | 0.11 | -10.24 | < 0.001 |
| DB | Low | 31 | 0.51 | 0.22 | -12.49 | < 0.001 | 0.45 | 0.22 | -13.83 | < 0.001 |
|  | Moderate | 31 | 0.55 | 0.21 | -11.86 | < 0.001 | 0.47 | 0.23 | -12.87 | < 0.001 |
|  | High | 31 | 0.60 | 0.17 | -12.86 | < 0.001 | 0.50 | 0.19 | -14.37 | < 0.001 |

AB, area-based, DB, distance-based, SSp, spatial summation of pain, SD, standard deviation

**Table 4.** Curve fitting: paired t-tests results for logarithmic and linear comparisons

| SSp type | Intensity | Mean difference | 95% CI | $t$ | $p$ |
| --- | --- | --- | --- | --- | --- |
| AB | Low | 0.05 | (0.02 - 0.08) | 3.46 | < 0.01 |
|  | Moderate | 0.06 | (0.04 - 0.09) | 4.95 | < 0.001 |
|  | High | 0.09 | (0.06 - 0.12) | 6.58 | < 0.001 |
| DB | Low | 0.06 | (0.03 - 0.09) | 3.97 | < 0.001 |
|  | Moderate | 0.08 | (0.05 - 0.11) | 5.11 | < 0.001 |
|  | High | 0.10 | (0.07 - 0.13) | 6.46 | < 0.001 |

AB, area-based, DB, distance-based spatial summation of pain, SSp, spatial summation of pain, CI, confidence interval. Note: Positive values in mean difference indicate better fit logarithmic curves.

General Linear Model performed on beta coefficients from logarithmic fittings showed a significant main effect for the factor ‘SSp’ (*F*_(1,30)_ = 164.66, *p* < 0.001, *η*_*p*_ = 0.85) and ‘intensity’ (*F*_(2,60)_ = 17.02, *p* < 0.001,*η*^2^_*p*_ = 0.36), indicating that i) in-general, area-based SSp was characterized by higher beta values and ii) that moderately painful (*p* < 0.001) and highly painful (*p* < 0.001) stimuli lead to a more dynamic pain increase, compared to the least painful stimuli (Fig. 5). No difference between moderately and highly painful stimuli was detected (*p* = 0.81, Fig. 5). Moreover, the GLM showed a significant ‘SSp’ *×* ‘intensity’ interaction (*F*_(2,60)_ = 5.18, *p* < 0.01, *η*^2^_*p*_ = 0.15) wherein post-hoc comparisons confirmed that: i) in the area-based paradigm, moderately (*p* < 0.01) and highly painful stimuli (*p* < 0.01) led to higher beta values than SSp induced via stimuli of the least painful intensity (Fig. 5), ii) distance-based SSp showed significantly higher beta-values for condition with high pain compared to the least painful stimuli, only (*p* < 0.05, Fig. 5). Non-linear fits showing that spatial summation of increases logarithmically are presented in Fig. 6 and Table 5.

**Table 5.** Descriptive statistics for logarhitmic and linear fits for SSp

| SSp type | Intensity | Logarithmic |  |  |  | Linear |  |  |  |
| --- | --- | --- | --- | --- | --- | --- | --- | --- | --- |
| | | $\beta$ | | $R^2$ | | $\beta$ | | $R^2$ | |
|  |  | Mean | SD | Mean | SD | Mean | SD | Mean | SD |
| AB | Low | 21.26 | 8.26 | 0.86 | 0.11 | 8.32 | 3.29 | 0.81 | 0.11 |
|  | Moderate | 27.98 | 9.64 | 0.86 | 0.09 | 10.74 | 3.68 | 0.80 | 0.11 |
|  | High | 26.15 | 9.61 | 0.88 | 0.07 | 9.93 | 3.65 | 0.79 | 0.11 |
| DB | Low | 11.52 | 6.96 | 0.51 | 0.22 | 4.19 | 2.91 | 0.45 | 0.22 |
|  | Moderate | 13.79 | 8.67 | 0.55 | 0.21 | 4.89 | 3.61 | 0.47 | 0.23 |
|  | High | 14.57 | 7.00 | 0.60 | 0.17 | 5.23 | 2.77 | 0.50 | 0.19 |

AB, area-based, DB, distance-based spatial summation of pain, SSp, spatial summation of pain, SD, standard deviation

**Figure 5.**
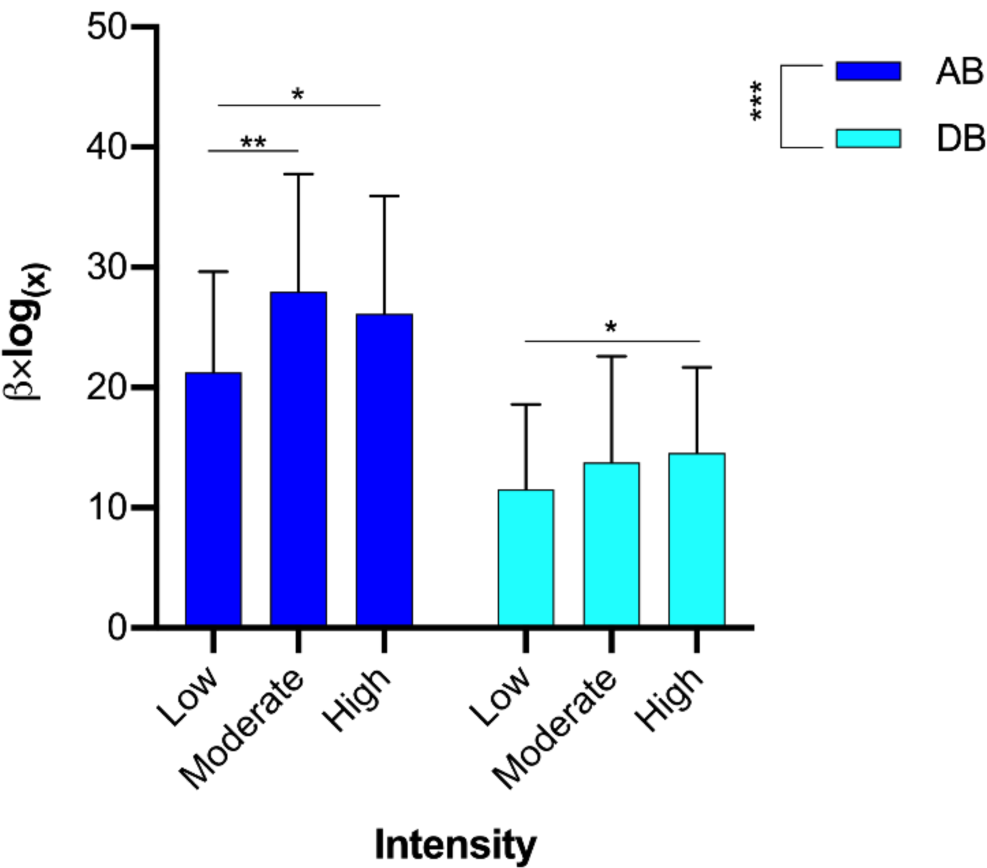
The magnitude of spatial summation of pain (SSp). Spatial summation of pain was expressed as beta coefficients of non-linear logarithmic functions. Higher beta values for area-based (AB) compared to distance-based (DB) were observed. The magnitude of SSp was pain intensity-dependent: when stimuli were calibrated to induce pain at the level of approximately 70 out of 100, the pain then increased more rapidly compared to pain at the level of 30. The figure presents mean values with standard errors of the mean (SE). \**p* < 0.05, \*\**p* < 0.01.

**Figure 6.**
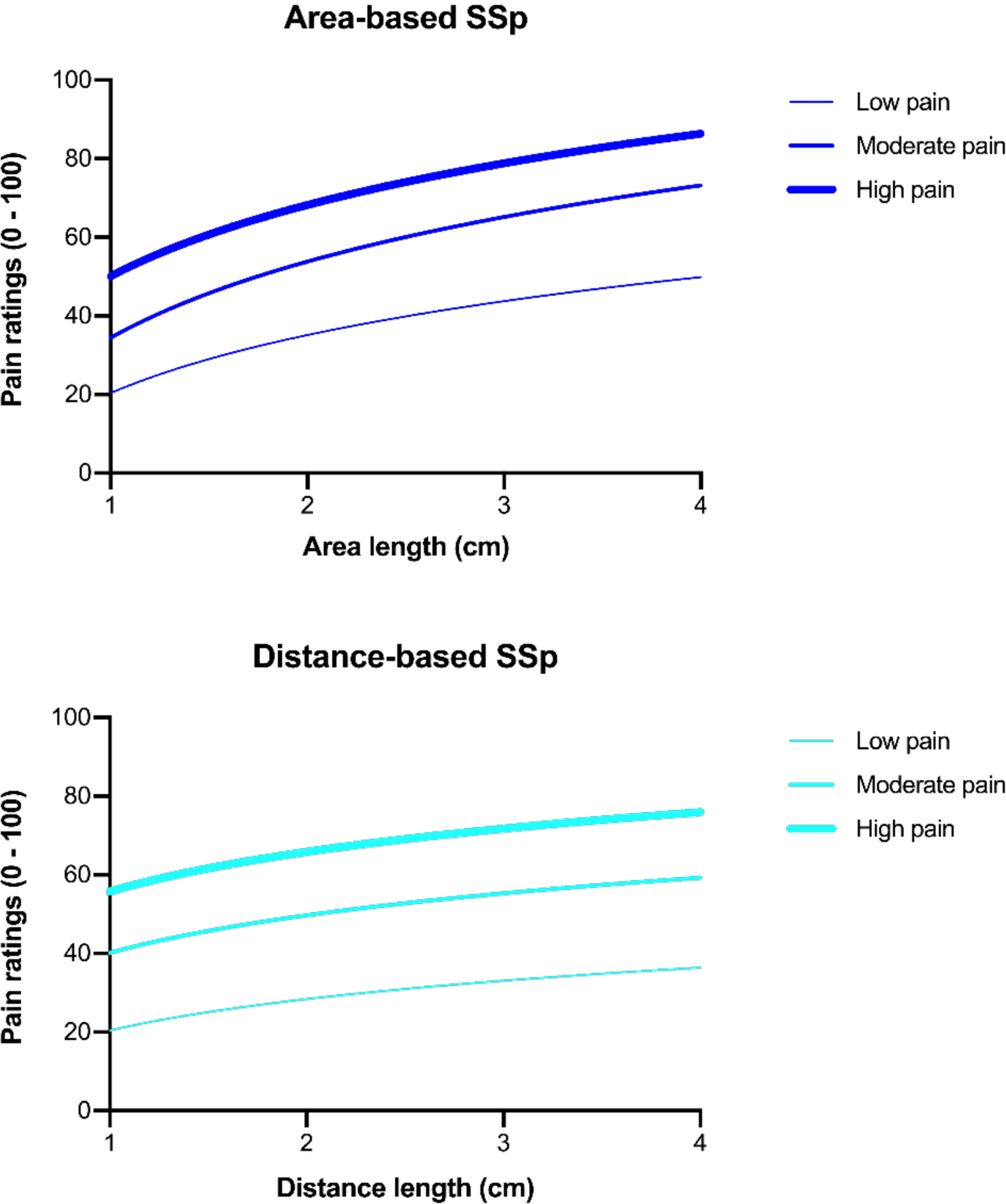
Spatial summation of pain increases logarithmically. The pain increases with increasing the area stimulated or the distance between two stimuli in a logarithmic fashion.

## DISCUSSION

The aim of this study was to investigate two types of SSp and to compare the summation trajectories when using individually calibrated noxious electrocutaneous stimuli. Three novel findings are reported, relating to the processing of the spatial information in the pain system: It was found that the pain increases logarithmically as a non-linear function and that the pattern of pain increase was statistically different across three different pain intensities, however in a not anticipated direction. SSp became saturated in the distance-based but not in the area-based SSp. Moreover, saturation was more likely to be observed when lower intensities were applied. Such an observation was possible to capture using curve fitting. The summation of pain was expressed as a function and decomposed to the value of one logarithmic coefficient. Coefficients were of higher values when the baseline pain level was higher than 50. And finally, data from our paradigm showed that area-based SSp is more painful than distance-based SSp, however this effect depends on how the painful stimuli are.

The long history of research on SSp encountered difficulties from the very beginning. For example, early reports failed to show SSp at all, possibly due to the insufficient stimuli intensity [18,21]. Later reports confirmed that stimulus intensity shapes the magnitude of SSp [38,42,54], suggesting that larger SSp is more likely to be observed in higher stimulation intensities. The putative mechanism of this relative increase in summation was attributed to a more pronounced excitation of spinal nociceptors at higher intensities [42]. These studies, however, have shown thermal SSp, expressed as the raw difference in pain ratings between two [42,54] or a maximum of three conditions (areas) [38]. The limited number of probes determined by the equipment used in previous studies, prevented from testing a predictive model. In the current study, five planar concentric electrodes were used for each SSp type, which allowed to investigate a summation pattern. It was found that a logarithmic function described more variance of the data compared to a linear fit. Interestingly, this pattern was observed for low, moderate and high pain conditions, confirming the validity of the model. The beta values were, however, higher when the intensity was calibrated to induce a high level of pain (>70/100), confirming preliminary observations, where lower SSp was observed for low pain intensities. This seems like a paradox. On the one hand, the pain increase is i) disproportional and ii) non-linear, but on the other hand, more intense stimuli elicit a larger SSp effect. The former results might indicate inhibitory processes underlying SSp, while the later points towards the contribution of larger excitation waves. In some circumstances, the non-linear nature of SSp might be of advantage for any living organism: Inhibitory processes saturate pain perception but not for very intense pain. Considering that the pain level may in acute situations reflect the magnitude of potential tissue damage, the bigger the damage (nociception) the bigger the pain increase. Thus, it seems that the greater magnitude of the SSp at higher intensities serves a protective mechanism preventing the individual from massive tissue damage.

The paradox described here may be explained by the variety of neural processes involved in the human [3,40] and non-human [6,22] neuroaxis, including descending pain inhibition [6], lateral inhibition [40] or even expectancies. Spatial summation has been originally linked with facilitation at the single neuron level. For example, Sherrington (1906) has used this term to describe the generation of action potential when subliminal -separated in space-stimuli were applied. Interestingly, first experiments on SSp in rats showed spatial summation to be disproportional [6]. Bouhassira et al. [6] found that convergent neurons exhibit enhanced electrical activity only when the spinal cord was sectioned and the effect of the descending loop was thereby reduced. This finding, although limited to rat studies, indicates that descending pain inhibition might play a role in the SSp magnitude and shape. Furthermore, a vast majority of human studies found similar observations: SSp has been reported as subadditive, as the reported pain marginally or modestly increased even though the stimulated area was doubled [10], tripled [41], quadrupled [54], or even six times larger [27]. The subadditivity supports the role of descending pain system. Indeed, the periaqueductal grey matter (PAG) has been found as core midbrain region of pain modulation [20,26,35]. Enhanced activity of this region has been found in hypoalgesic or hyperalgesic modulation, which presumably is reflected in its anatomical structure [30]. Whether PAG activation is intensity-dependent is unclear and warrants future investigations.

The fact that a significant interaction between the type of SSp and the relative painfulness was detected indicates that, potentially, lateral inhibition interplayed with SSp. For example, in the study by Quevedo et al. [40], those trials were much more painful, in which only two points were separated by e.g. 8cm compared to a line (8cm length) stimulus applied by laser. In the current study, when the high intensity condition was applied, there was no difference between area- and distance-based SSp which is partially in line with this previous report. Although this current data did not reveal a similar trend as in the study by Quevedo et al. [40], an interesting observation was, that despite a smaller number of nociceptors activated in distanced-based SSp, an equal pain responses were noticed. The lack of replication of the full findings of the previous report are methodological differences, mostly related to stimuli types and timing of stimuli application. In that sense, for an inhibitory response, the temporal relation between spatially distinct stimuli might be of important for the prediction of an accurate SSp effect.

For the first time, area-based and distance-based SSp were compared in terms of the pattern of pain summation. Area-based SSp was not saturated over the course of the linear increase in size of the stimulated area. Pain successfully increased, when the area increased but not when the distance between two points became larger. Furthermore, this is the first report which proved that SSp is a local effect as we used a continuum of areas, or distances, in a line-like manner of up to 4cm, therefore the current findings complement previous SSp research using laser, cold, pressure or electrocutaneous stimuli at larger areas.

Although the current findings are robust in terms of internal validity, some aspects could have affected the SSp estimations: planar concentric electrodes were placed close to each other with no space in between. This solution was sufficient to manipulate area size; however, they were connected via only one tangential point. Such a solution has been used previously, yet for future experiment it is recommended to use square-like stimuli. Trains of pulses produced a strong pain at a relatively low intensity (<12 mA). This could be a result of temporal summation processed as ‘noise’ during the spatial summation. Notwithstanding to that, temporal facilitation was limited to single-stimulus only, and seems unlikely to affect the pattern of spatial summation described in current study. The paradigm used in this study was based on a random sequence of different stimulus types, with three repetitions each. Such a small number of trials however, was compensated by three separate examination sessions for each participant, which likely affected the precision of SSp estimations. Furthermore, as no difference between the three sessions was detected it seems that the paradigm used and SSP effect itself is stable over-time.

### Conclusions

Taken together, results presented here are an important step forward in the understanding of the pain prediction in relation to spatial configurations of noxious stimuli. Our data indicates that the relative “painfulness” of noxious stimuli affects the non-linear summation trajectory, however, the hypothesis that reduced SSp magnitude is observed when high intensity of stimuli is used was not supported. It seems however, that SSp increases logarithmically, which may suggest the SSp effect is unique mixture of facilitatory and inhibitory processes. Current data provide support for the lateral inhibition in the nociceptive system as under some circumstances the same pain level was reported in both SSp types even though one of them involved a greater area of nociceptive stimulation.

## ACKNOWLEDGEMENTS

Authors declare no conflicts of interest. Data presented here constitute a part of the preregistered project (https://osf.io/qry9d) dedicated to test if the pain intensity affects magnitude and reliability of SSp. Due to the significant amount of data generated in this project, reliability results will be presented in other occasion.

## Footnotes

1 Semantically, a more correct way to describe SSp in animal is spatial summation of nociceptive processing. This is due to the IASP definition of pain and its recent updated version, according to which, pain is a subjective experience.

## Notes

https://osf.io/qry9d

### Competing Interest Statement

The authors have declared no competing interest.

### Summary of Updates

Some technical details have been added. Language has been corrected, spelling and types were corrected.

